# Anaerobe isolation from denitrifying benzene-degrading enrichment culture and their capacity to mineralize benzene

**DOI:** 10.1101/2023.01.07.522375

**Authors:** Samuel C Eziuzor, Carsten Vogt

## Abstract

Only a few benzene-mineralizing anaerobes have been isolated to date. In an attempt using classical isolation techniques to isolate benzene-mineralizing pure cultures from a benzene-mineralizing nitrate-reducing microbial community, two putative isolates were gained under nitrate-reducing conditions spiked separately with acetate and benzene as sole sources of carbon and energy with media containing ammonium or without ammonium. Both putative isolates; Bz4 (with ammonium) and Bz7 (without ammonium) - mineralized ^13^C-labelled acetate under anoxic conditions at 3.3 and 2.7 μM day^-1^, respectively, revealed by analysis of evolved ^13^CO_2_. However, only Bz4 mineralized [^13^C_6_]-labelled benzene (0.298 μM benzene mineralized day^-1^) generated up to 960.2 ± 0.3 ‰ δ^13^C-CO_2_ during 184 days while producing only slight amounts of nitrite (4.60 ± 0.004 μM); no benzene was mineralized by Bz7 during 184 d, and no nitrite was detected. The 16S rRNA gene amplicon sequencing of the acetate-grown bacteria revealed consortia enriched in *Nocardioides* (8.9%), *Pseudomonas* (18.2%), *Rhizobiaceae* (21.0%), *Allorhizobium-Neorhizobium-Pararhizobium-Rhizobium* (51.4%) for Bz4 and S*implicispira* (96.7%) for Bz7. The gained Bz4 consortium that mineralized benzene under anoxic condition can be further purified and explored for their metabolic potentials.

## 1. Introduction

Several isolation attempts demonstrated that the majority of bacteria cannot be cultured in the laboratory yet (Rappe and Giovannoni, 2003). The isolation of anaerobic benzene degrading bacteria directly from enrichment cultures spiked with benzene, over the years, has been largely unsuccessful. However, very few pure cultures of anaerobes capable of oxidizing benzene have been isolated under nitrate-and iron-reducing conditions (Coates et al., 2001; Kasai et al., 2006; Duo et al., 2010; Holmes et al., 2012; Zhang et al., 2012; Devandera et al., 2019). Successful isolates using nitrate as an electron acceptor and capable of benzene degradation include *Azoarcus* species isolated from gasoline-contaminated groundwater in Kumamoto, Japan (Kasai et al., 2006). The genome of the benzene-degrading *Azoarcus* strain DN11 was later published by Devanadera et al. (2019). In addition, a strain of the species *Bacillus cereus* growing on benzene under nitrate-reducing was isolated by screening procedure from gasoline-contaminated soil (Dou et al., 2010). A hyperthermophilic archaeon, *Ferroglobus placidus* is capable of benzene mineralization coupled to the reduction of Fe(lll) under strictly anoxic conditions, demonstrated by applying ^14^C-labelled benzene as sole organic carbon substrate and subsequent analysis of produced ^14^CO_2_ (Holmes et al., 2011). *Geobacter mettalireducens* and strain Ben belonging to the genus *Geobacter*, were reported to grew in medium with benzene as the sole electron donor and Fe(III) oxide as the sole electron acceptor (Zhang et al., 2012). So far, no isolate has been reported to mineralize benzene under sulfate-reducing or methanogenic conditions.

The focus of this study is on nitrate as a common electron acceptor in aquifers due to the usage of nitrate-based fertilizers, resulting in widespread global groundwater contamination by nitrate. In addition, nitrate is an energetically favourable electron acceptor compared to sulfate or carbonate, which is also used by several organisms. Dissimilatory nitrate reduction is one of the key processes in nitrate reduction which can lead to nitrate reduction to dinitrogen gas or dissimilatory nitrate reduction to ammonium, both are used for respiration (Kuypers et al., 2018). Notably, several nitrate-reducing aromatics degrading pure cultures have been isolated, which makes it promising to try to isolate benzene degraders at nitrate-reducing conditions (Weelink et al., 2010). Here, we gained two putative isolates dominated by *Gammaproteobacteria* that were obtained by classical isolation techniques before incubation with ^13^C-benzene to assess their capacity to mineralize benzene under denitrifying conditions and characterized by analysis of 16S rRNA gene sequence.

## 2: Materials and methods

### 2.1: Chemicals

Chemicals were purchased from Fluka (Steinheim, Germany), Merck (Darmstadt, Germany), Roth (Karlsruhe, Germany), and Sigma-Aldrich (Taufkirchen, Germany) in p.a. quality if not otherwise stated. Benzene-^13^C_6_ and acetate-^13^C_2_ were purchased from Sigma-Aldrich with an isotopic purity of 99 atom % ^13^C.

### 2.2: Isolation procedures

A denitrifying benzene-mineralizing culture enriched from a former benzene-contaminated aquifer near Zeitz in Germany in *on-site* columns containing coarse sands; originally enriched as a sulfate-reducing culture but later was adapted to the denitrifying condition and maintained for several years in the laboratory (Vogt et al., 2007; Keller et al., 2018; Eziuzor et al., 2022). The enrichment culture is made up of both liquid-phase and solid phases comprising coarse sands. A hundred microliters of the liquid phase from twice inverted microcosm were spread-plated to initiate isolation on agar plates containing 8 mM sodium nitrate for approaches with ammonium (7.5 mM) and without ammonium in the anaerobic glove box. Mineral salt medium-agar (MSM-agar) was solidified with 1.5% granulated agar-agar (Merck, Darmstadt, Germany). Plates were stacked in an anaerobic jar and anaerobic conditions were generated using AnaeroGen™ 2.5L (ThermoScientific, Oxoid Ltd, UK). Benzene was provided via vapour-phase transfer by the headspace of the anaerobic jar, using a filter paper (70 mm Ø filter paper; LLG Labware, France) impregnated with 20 μl pure benzene. Live and abiotic controls were setups by incubating inoculated agar plates without providing benzene via vapour-phase transfer in the headspace as well as abiotic controls by providing non-inoculated agar plates with vapour-phase benzene transfer for both approaches with ammonium and without ammonium. The setup was incubated at room temperature in the dark until visible colonies were formed. Colonies were streak-plated onto new MSM-agar plates aseptically in the anaerobic glove box. Streak plates were incubated in the anaerobic jar as described above until discrete colonies were visible. The discrete colonies were carefully picked and suspended into a 30 mL nitrate-amended mineral salt medium in 50 mL serum bottles (Glasgerätebau Ochs, Bovenden-Lenglern, Germany). Serum bottles were sealed with gas-tight inert Teflon-coated butyl rubber stoppers (ESWE Analysentechnik, Gera, Germany) and spiked with 0.05 mM benzene using a sterile 10 μL microlitre gas-tight syringes (Hamilton, Switzerland). The microcosms were incubated statically at room temperature in the dark for possible growth. Growth in the microcosms was checked after nine months through cell count analysis by staining with SYBER-Green I (Thermo Fischer Scientific GmbH, Germany) and visualized with an Epifluorescence microscope (Nikon, Minato City, Japan). All plating and transfers were set up in an anaerobic glove box (Coy Laboratory Products Inc., Grass Lake, USA) containing a gas atmosphere of 95% N_2_ and 5% H_2_. The schematic representation of approaches adopted in attempting to isolate putative denitrifying benzene-mineralizing anaerobes from the benzene-enrichment community is shown in Figure 1.

**Figure 1.**
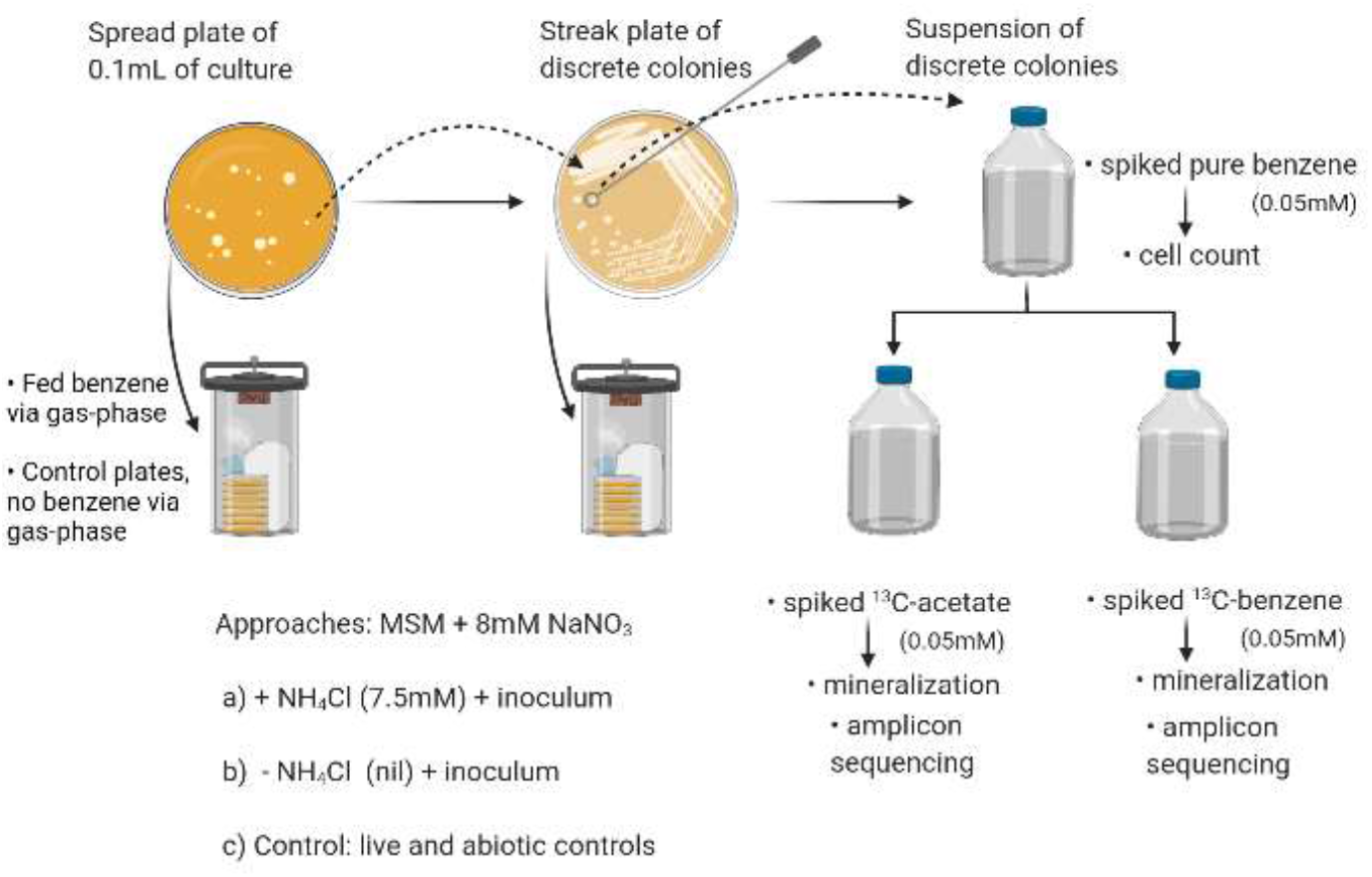
Schematic representation of approaches adopted in attempting to isolate putative benzene-mineralizing anaerobes from a benzene-degrading nitrate-reducing community. Formed colonies were suspended in a nitrate-containing anoxic mineral salt medium (with or without ammonium) spiked with 0.05 mM of ^13^C-acetate or ^13^C-benzene to check for mineralization. Cultural setups were incubated at room temperature in the dark. The figure was created with Biorender.

### 2.3: Mineralization and nitrite analyses

Ten percent transfer was made into a 30 mL MSM in 50 mL serum bottles and spiked with either ^13^C-acetate or ^13^C-benzene at the concentration of 0.05 mM to ascertain growth and mineralization. Transfers were made in MSM with ammonium or without ammonium in addition to abiotic controls. Headspace samples were taken by sterile syringes previously flushed with nitrogen to exclude oxygen and the carbon isotope ratio of produced CO_2_ was determined using a gas chromatograph-isotope ratio mass spectrometer (GC-IRMS) as described elsewhere (Herrmann et al., 2010; Eziuzor et al., 2021). Carbon isotope ratios were expressed in the delta notation in per mil (d ^13^C/‰) units relative to the Vienna Pee Dee Belemite (VPDB) according to Coplen (2011). Liquid samples for nitrite analyses were taken equally as described above. The concentrations of nitrite were spectrophotometrically determined according to Raihan and colleagues (1997). Precisely, 125 μL of nitrite determination reagent (10 g/L sulfanilamide + 0.5 g/L napthylethylenediamine dihydrochloride (NEDD) dissolved in 10% H_3_PO_4_) was added to 500 μL sample and incubated after vortexing for 10 min in the dark. Absorbance was measured at 540 nm against a standard consisting of 125 μL determination reagent mixed with 500 μL distilled water. Quantification was done with external standard calibration.

### 2.4: Genomic DNA extraction and 16S rRNA gene sequencing

Genomic DNA was extracted from only the ^13^C-acetate grown cultures that yielded enough DNA at 94 days of incubation using the NucleoSpin Microbial DNA kit (Macherey-Nagel) according to the manufacturer’s instructions. The quality and quantification of extracted DNA were done using a NanoDrop ND 1000 spectral 183 photometer (Thermo Fisher Scientific, United States) and a Qubit fluorometer using the Qubit dsDNA BR assay kit (Thermo Fisher Scientific GmbH, Germany). The extracted DNA was stored at -20°C until further use. The extracted genomic DNA were PCR-amplified using the primers of 16S rRNA genes according to Klindworth et al. (2013). Amplicon sequencing was carried out using an Illumina MiSeq platform (Illumina, CA, USA) generating 2×250 bp paired-end reads. Sequencing reads were trimmed of primers and amplicon sequence variants (ASVs) were generated using DADA2 (Callahan et al., 2016). Taxonomic assignment was done using Silva version 132 (Bolyen et al., 2019) as a reference database through QIIME 2 (Quast et al., 2013). The demultiplexed raw reads of amplicon sequencing are available under the accession numbers ERR9078369 and ERR9078370 as submitted to the European Nucleotide Archive (ENA; www.ebi.ac.uk/ena).

## 3: Results

The picking of discrete colonies formed on agar surfaces was done based on their colony morphologies (whitish or brownish, circular raised or flat). In total, 8 colonies: 4 colonies each were obtained from MSM-agar plates with ammonium or without ammonium. Colonies were also observed on the live control incubated without benzene in the headspace. There was no growth observed in the abiotic controls. The suspended colonies in nitrate-containing anoxic mineral salt medium spiked with non-labelled benzene was checked for bacterial growth after 190 days and only two out of the 8 transfers are reported in Figure 2.

**Figure 2.**
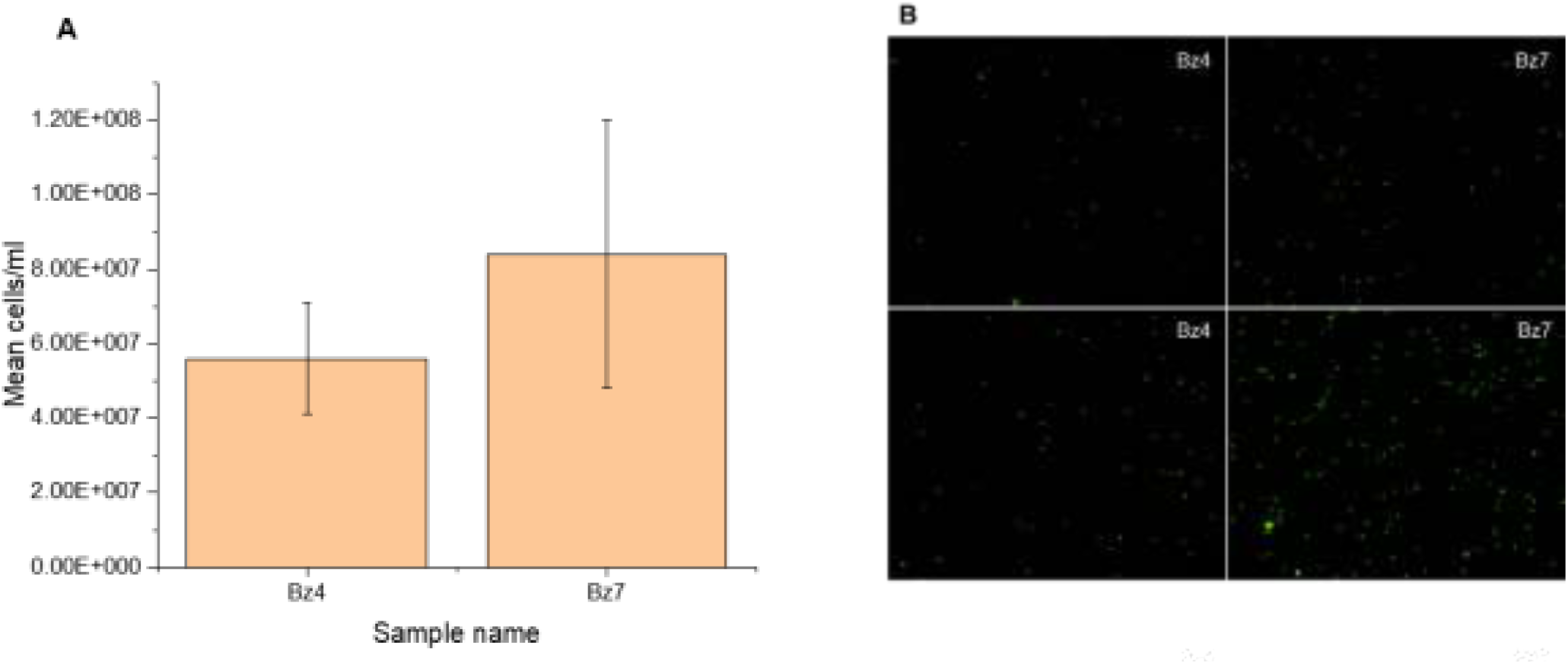
Cell counts (a) and photomicrograph of stained cells (b) using SYBER Green stain visualized with an Epifluorescence microscope of the transferred putative discrete colonies from streak plates after 190 days of anaerobic incubation in liquid MSM medium spiked with benzene and nitrate. The counted volume was 0.00117 μl with an image width of 0.18037 μm and an image height of 0.14376 μm per view.

Subsequently, one transfer each from approaches with ammonium (Bz4) and without ammonium (Bz7) is presented here based on their mineralization and 16S rRNA amplicon sequencing results. Mineralization analysis revealed that both putative isolates – Bz4 and Bz7 mineralized acetate under anoxic conditions at 3.3 and 2.7 μM day^-1^, respectively (Table 1).

**Table 1.** Mineralization of ^13^C-labelled acetate and benzene to ^13^C-CO_2_ under denitrifying conditions and nitrite concentrations at 94 days and 184 days, respectively for the two consortia.

| *Consortia* | *Acetate-grown* |  |  | *Benzene-grown* |  |  |
| --- | --- | --- | --- | --- | --- | --- |
| | $\delta^{13}\text{C}$-CO<sub>2</sub><br>(‰) | Mineralization (μM<br>acetate day<sup>-1</sup>) | Nitrite (μM) | $\delta^{13}\text{C}$-CO<sub>2</sub><br>(‰) | Mineralization (μM<br>benzene day<sup>-1</sup>) | Nitrite (μM) |
| *Bz4* | 5556.09±4.09 | 3.302 | 1.00±0.002 | 960.15±0.31 | 0.298 | 4.60±0.004 |
| *Bz7* | 4537.11±2.01 | 2.702 | 0.20±0.001 | -15.29±0.15 | 0.000 | 0.30±0.002 |

Each data point for $^{13}\text{C}$ -CO<sub>2</sub> and nitrite is the average of all replicate cultures.

However, only Bz4 could mineralize benzene under the incubation conditions and be considerably enriched in δ^13^C up to 960.2 ± 0.30 ‰ δ^13^C-CO_2_ on 184 d while no benzene was mineralized in Bz7 after 184 d (Figure 3).

**Figure 3.**
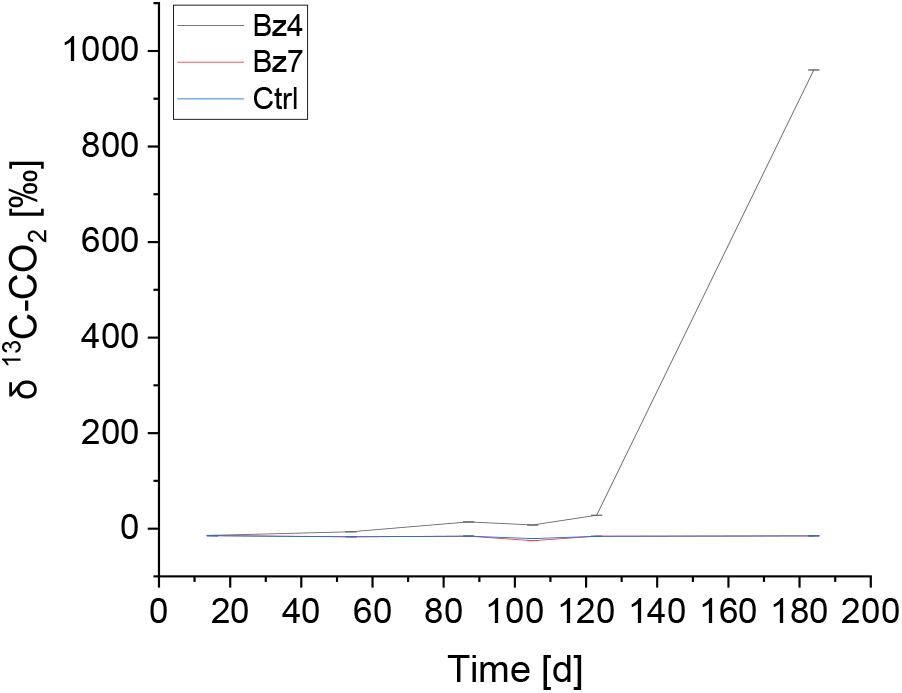
Mineralization of benzene under denitrifying conditions by putative isolates from the medium supplemented with ammonium chloride (Bz4), and without ammonium chloride (Bz7) in the medium. Each point was measured in triplicates with standard deviation.

The nitrite concentration was not regularly measured due to the small volume of the culture medium, however, it was measured in triplicates only at the last sampling (Table 1). Nitrite concentrations of 1±0.002 μM and 4.6 ± 0.004 μM were observed for the acetate-grown and benzene-grown Bz4 at 94 d and 184 d, respectively while no nitrite concentration increases were observed in acetate-grown consortia at 94 d in spite of acetate mineralization.

The 16S rRNA gene amplicon sequencing of the acetate-grown consortia revealed that the putative isolates were not pure but rather consortia enriched in *Nocardioides* (8.9%), *Pseudomonas* (18.2%), unknown *Rhizobiaceae* (21.0%), and *Allorhizobium-Neorhizobium-Pararhizobium-Rhizobium* (51.4%) for Bz4 while uncultured *Anaerolineaceae* (1.2%) and *Simplicispira* (96.7%) for Bz7 as shown in Figure 4. The benzene-grown consortia did not yield enough biomass for DNA extraction after 184 days of incubation.

**Figure 4.**
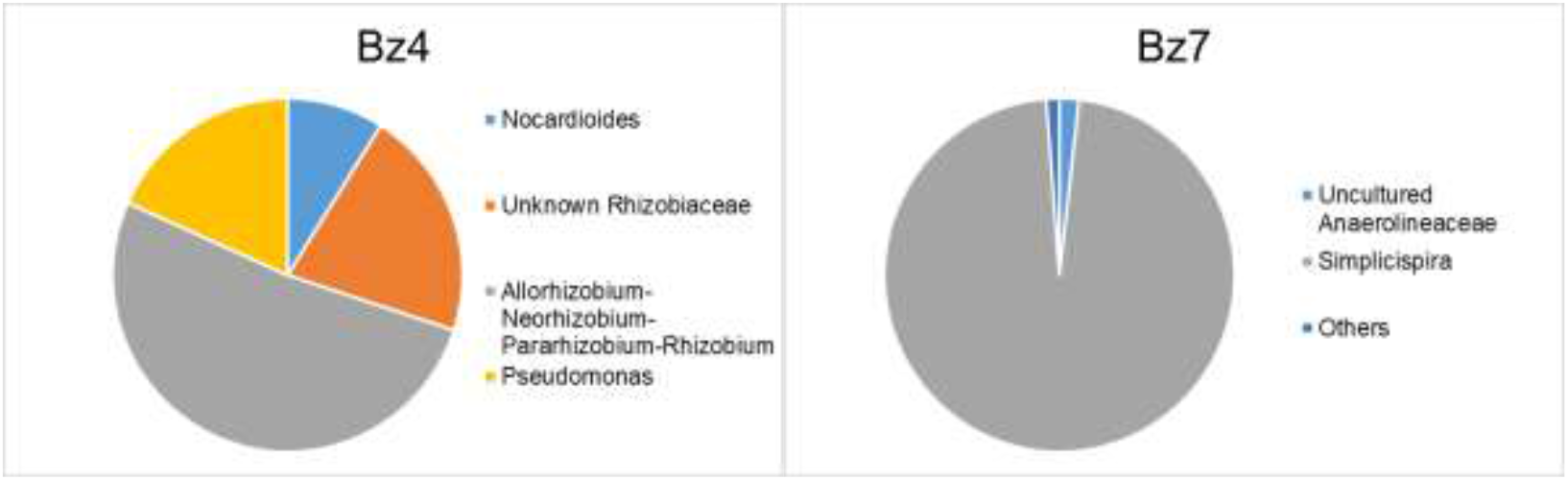
16S rRNA gene amplicon sequencing of the putative isolates on day 94 for ^13^C-acetate-grown consortia at the genus level. Bz4 was cultured with ammonium while Bz7 was cultured without ammonium.

## 4: Discussion

Bacterial growth on the benzene-fed MSM-agar plates, in the presence or absence of ammonium, and on the live control resulted in distinct colony morphologies, indicating that they support growth. The observed colonies on the live control are due to some organics in the MSM-agar, which supported growth. The absence of observable growth on the abiotic controls showed that the observed colonies on the live approaches emanated completely from the original inoculum. The cell counts - mean cell per mililitre 5.60E+07 to 8.4E+07- and photomicrograph showed that the liquid cultures contained a distinct cell number after longer incubation (190 days), indicating that cells grew with benzene. Anaerobes are known to have a long lag phase and, in this case, would require more time to acclimatize in the medium. A lag phase of around 70 days was observed in the original benzene-mineralizing, nitrate-reducing microcosms (Eziuzor et al., 2022). Considering the long lag phases observed in anaerobic cultivation and the inherent acclimatization of the cells in their new, unnatural media, it is expected that the isolates will take even a longer time to acclimatize, proliferate and metabolically degrade the stable benzene compound only if favoured anaerobically.

The consortium Bz4 was capable to mineralize ^13^C-benzene up to 0.298 μM benzene day^-1^ in the presence of nitrate as a terminal electron acceptor as the nitrite content increased to 4.6 μM. The consortium was cultured with ammonium in the medium indicating that the denitrification process was operative. Acetate was easily mineralized by both consortia as it serves as a common intermediate in several metabolic processes (Lueders, 2017) though no nitrite increase was detected in Bz7 cultivated without ammonium. The nitrite concentration was higher in Bz4 compared to Bz7 and higher in benzene-grown strains than the acetate-grown strains. Unfortunately, nitrate was not measured to prove its usage as an electron acceptor. The nitrite production as observed indicated mineralization coupled with nitrate reduction was active in Bz4 unlike in Bz7 (Table 1). This could be due to the presence of ammonium in the medium for Bz4 which supported the nitrate reduction process. Nitrate reducers are known to assimilate ammonium for their growth, however, in the absence of ammonium in the medium, certain microbes are known to produce ammonium (ammonification) from nitrate and other medium components. The absence of ammonium in the medium was to investigate if denitrification will proceed while producing ammonium required for growth as the absence of ammonium prevents the growth of organisms performing Anammox. Usually, nitrate is used as an electron acceptor or used as a nitrogen source; in the latter case, the formed ammonium is incorporated in biomolecules (e.g. amino acids, proteins). However, if nitrate is used as an electron acceptor by ammonification, the formed ammonium can be used by other organisms as a nitrogen source or used by Anammox organisms as an electron source.

All bacterial phylotypes identified in the acetate-grown consortia as uncultured *Anaerolineaceae* (0 – 4.4%) (Family: *Anaerolineaceae* (1 – 9%)); *Simplicispira* (0 – 3.2%) (Family: *Burkholderiaceae* (2 – 11%)); *Nocardioides* (0 – 6.9%) (Family: *Nocardioidaceae* (0 – 9%)); *Pseudomonas* (0 – 11.4%) (Family: *Pseudomonadaceae* (0 – 21%)), and unknown *Rhizobiaceae* (0 – 0.2%), *Allorhizobium-Neorhizobium-Pararhizobium-Rhizobium* (0 – 0.2%) (Family: *Rhizobiaceae* (< 1%)) were detectable at the indicated range of relative abundances throughout the time-course of the previous study done with the original inoculum (Eziuzor et al., 2022). Thus, the corresponding phylotypes identified in acetate-grown consortia showed a snapshot of possible isolates under the classical isolation procedure employed here. Notably, the key player for primary benzene degradation postulated in the previous study, the distinct strain of the *Peptococcaceae* was not enriched among others like *Brocadiaceae* (1 – 12%), *Ignavibacteriaceae* (5 – 42%) and *Rhodocyclaceae* (7 – 29%), which have higher relative abundances in the original culture (Eziuzor et al., 2022). We posit that it is rather unlikely that the *Peptococcaceae* was enriched here because the organism may not use acetate at all as a carbon source even with nitrate as an electron acceptor.

Consortium Bz7 maintained without ammonium was highly enriched in *Simplicispira* at 96.7% relative abundance. *Simplicispira* is a Gram-negative, motile and facultative anaerobic rod-shaped denitrifying bacteria (Siddiqi et al., 2020). Species of the genus *Simplicispira* contain ubiquinone-8 (Q-8) as a major quinone with a DNA G + C content of 63–65 mol% (Grabovich et al., 2006). The genus *Simplicispira* so far comprises six described species with validly published names: *Simplicispira metamorpha, Simplicispira psychrophila* (Grabovich et al., 2006), *Simplicispira limi* (Lu et al., 2007), *Simplicispira piscis* (Hyun et al., 2015), *Simplicispira suum* (Cho et al., 2018), and *Simplicispira hankyongi* (Siddiqi et al., 2020). Bacosa et al. (2021) reported that *Burkholderia* (B5) preferentially degraded benzene over octane, an aliphatic hydrocarbon, with varying concentrations of up to 4.35 mM. B5 was found to possess alkane hydroxylase (alkB) and catechol 2,3-dioxygenase (C23D) genes, which are responsible for the degradation of alkanes and aromatic hydrocarbons, respectively in the presence of oxygen. Some members of the *Burkholderiales* were classified as benzene consumers in a benzene-degrading nitrate-reducing culture (Melkonian et al., 2021) and their potential to make enzymes for the degradation of benzoate and hydroxybenzene, two intermediates of anaerobic benzene degradation, reported (van der Waals et al., 2017). Interestingly, genes encoding enzymes for benzene degradation that typically combine anaerobic steps with aerobic ones, one of which includes oxygenation to convert benzoyl-CoA to acetyl-CoA and succinyl-CoA were expressed in their culture (Melkonian et al., 2021). This may seem surprising for an anoxic culture (Melkonian et al., 2021), and might be the situation in the Bz7 consortium if ammonium is provided. *Simplicispira* enriched in the Bz7 consortium did not mineralize benzene after 184 days of anaerobic incubation (Figure 3). This observation indicated that the taxon was incapable of anaerobic benzene oxidation in the incubation conditions employed which might be due to the absence of ammonium in the medium. It likely existed in the original consortium as a secondary products feeder. *Anaerolineaceae* are usually found to be associated the with anaerobic benzene degradation process under nitrate-reducing conditions (Atashgahi et al., 2018; Eziuzor et al., 2022). The consortium Bz7 has a 1.2% relative abundance of uncultured *Anaerolineaceae* which might be contributing to benzene-mineralization at nitrate-reducing conditions.

We assume that the presence of *Pseudomonas* (18.2 % relative abundance) as identified in the acetate-grown consortium Bz4 might be responsible for the benzene mineralization in the corresponding benzene-grown consortium (Figure 3). *Pseudomonas* comprise a genus-level group of facultative anaerobic bacteria within the class *Gammaproteobacteria*, notable for their metabolic diversity including the ability to degrade a variety of aromatic compounds. *Gammaproteobacteria* shows a close relationship to the *Betaproteobacteria*, which was drastically rearranged as an order of *Gammaproteobacteria* recently (Quast et al., 2013). Taxonomic analysis based on 16S rRNA gene sequences placed *Gammaproteobacteria* distantly to the other three classes of proteobacteria (*Alphaproteobacteria, Deltaproteobacteria* and *Epsilonproteobacteria*) (Gao et al., 2009; Gupta, 2000). There are several species of isolated *Pseudomonas* degrading benzene (Mahendran et al., 2006), *P. veronii* (1YdBTEX2 and 1YB2) (Lima-Morales et al., 2013; 2016), and toluene (1YdBTEX2) (Morales et al., 2016) in the presence of oxygen. They are capable of anaerobic growth in the presence of nitrate, nitrite, and nitrous oxide (N_2_O) as terminal electron acceptors (Wu et al., 2005). Thus, anaerobically several species have been studied such as *P. aeruginosa* for their proteome (Wu et al., 2005), *P*. sp. JP1 as high molecular weight PAH degrader (Liang et al., 2014), and *P. aeruginosa* strain PAH-1 as phenanthrene degrader (Ma et al., 2011). The mechanism of their metabolic degradation of aromatics under anoxic conditions is not clearly known.

Rhizobial genera of unknown *Rhizobiaceae* and *Allorhizobium-Neorhizobium-Pararhizobium-Rhizobium*, ANPR at various relative abundances in the consortia were identified (Figure 4). They are known nitrogen-fixing symbionts mostly found naturally in legumes (Mousavi et al., 2014) and have been identified in anaerobic microbial communities degrading acetate under sulfate-reducing conditions (Bin Hudari et al., 2020), and benzene under nitrate-reducing conditions (Eziuzor et al., 2022), which is the original inoculum for the isolation attempt. ANPR was also found to be the dominant phenanthrene degrader aerobically in gradient anthropized soils (Lemmel et al., 2019). They are likely found in the examples stated above as nitrogen-fixing opportunists due to the presence of nitrate reducers, thereby, contributing to the dissimilatory inorganic nitrogen metabolism

Interestingly, the isolation procedure did not result in the enrichment of the putative benzene degrader, *Peptococcaceae* phylotypes, which was identified in the original inoculum. This might be because *Peptococcaceae* and other phylotypes like *Ignavibacteriaceae* and *Rhodocyclaceae* cannot be enriched through the classical isolation technique employed here and may need to be selectively enriched in a liquid medium only. In addition, the identified organisms were pre-grown with acetate and not benzene before sequencing. The *Peptococcaceae* has been found to disappear even when benzene mineralization stopped in a time-course experiment coupled with nitrate reduction (Eziuzor et al., 2022). Melkonian et al. (2021) demonstrated that most of the benzene-degrading nitrate-reducing community members likely feed on metabolic left-overs or on necromass while only a few of them, from families *Rhodocyclaceae* and *Peptococcaceae*, are primary benzene degraders. The microbial consortia identified in Bz4 and Bz7 consortia likely have their metabolic functions in the original anaerobic microbial ecosystem under nitrate-reducing conditions as benzene-degrader, benzene-derived intermediates consumers, necromass, nitrate reducers, and nitrogen-fixing symbionts. The symbionts may have established mutualistic or syntrophic interactions with the *Gammaproteobcateria* leading to their living together though their interactions are not clear. Here, we gained a consortium, Bz4 which has the capacity to mineralize benzene under anoxic conditions, though not truly pure but can be further purified and explored for its metabolic potentials. However, this may also separate the mutualistic relationship of phylotypes. Alternatively, substrate enrichment can encourage the growth of the preferred candidate in order to preserve the difficult-to-grow organisms in the growth medium. Bz7 consortium might mineralize benzene if ammonium is provided in the culture medium to encourage the denitrification process.

